# Expanded genomic analyses for male voice-breaking highlights a shared phenotypic and genetic basis between puberty timing and hair colour

**DOI:** 10.1101/483933

**Authors:** Ben Hollis, Felix R. Day, Alexander S. Busch, Deborah J. Thompson, Ana Luiza G. Soares, Paul R.H.J. Timmers, Alex Kwong, Doug F. Easton, Peter K. Joshi, Nicholas J. Timpson, The PRACTICAL Consortium, 23andMe Research Team, Ken K. Ong, John R.B. Perry

**Affiliations:** MRC Epidemiology Unit, Institute of Metabolic Science, University of Cambridge School of Clinical Medicine, Cambridge Biomedical Campus Box 285, Cambridge, CB2 0QQ, UK; Department of Growth and Reproduction, Rigshospitalet, University of Copenhagen 2100 Copenhagen O, Denmark; International Center for Research and Research Training in Endocrine Disruption of Male Reproduction and Child Health, Rigshospitalet, University of Copenhagen, 2100 Copenhagen O, Denmark; Centre for Cancer Genetic Epidemiology, Department of Public Health and Primary Care, University of Cambridge, Cambridge, UK.; MRC Integrative Epidemiology Unit at the University of Bristol, Bristol, UK; Population Health Science, Bristol Medical School, University of Bristol, Bristol, UK; Usher Institute for Population Health Sciences and Informatics, University of Edinburgh, Teviot Place, Edinburgh, EH8 9AG, UK; IUMSP, Biopôle, Secteur Vennes-Bâtiment SV-A, Route de la Corniche 10, 1010 Lausanne, Switzerland; School of Geographical Sciences, University of Bristol, UK; Centre for Multilevel Modelling, University of Bristol, UK; 23andMe, Inc., Mountain View, California 94041, USA; Department of Paediatrics, University of Cambridge School of Clinical Medicine, Cambridge Biomedical Campus Box 181, Cambridge, CB2 0QQ, UK

## Abstract

The timing of puberty is highly variable and has important consequences for long-term health. Most of our understanding of the genetic control of puberty timing is based on studies in women, as age at menarche is often recorded. Here, we report a multi-trait genome-wide association study for male puberty timing, based on recalled timing of voice breaking and facial hair, with an effective sample size of 205,354 men, nearly four-fold larger than previously reported. We identify 78 independent signals for male puberty timing, including 29 signals not previously associated with puberty in either sex. Novel mechanisms include an unexpected phenotypic and genetic link between puberty timing and natural hair colour, possibly reflecting common effects of pituitary hormones on puberty and pigmentation. Earlier male puberty timing is genetically correlated with several adverse health outcomes and, in Mendelian randomization analyses, shows causal relationships with higher risk of prostate cancer and shorter lifespan. These findings highlight the relationships between puberty timing and later health outcomes, and demonstrate the value of genetic studies of puberty timing in both sexes.

## Introduction

The timing of puberty varies widely in populations and is determined by a broad range of environmental and genetic factors^1^. Understanding the biological mechanisms underlying such variation is an important step towards understanding why earlier puberty timing is consistently associated with higher risks for a range of later life diseases, including several cancers, cardiovascular disease and Type 2 diabetes^2–4^.

Most of our understanding of the genetic determinants of puberty timing is based on studies in women, as the age of first menstrual bleeding (age at menarche, AAM) is a well-recalled and widely measured marker of female sexual development. The largest genome-wide association study (GWAS) to date for AAM, in ~370,000 women, identified 389 independent signals, accounting for approximately one quarter of the estimated heritability for the trait^5^. In contrast, genetic studies of puberty timing in men are much fewer and smaller in scale, due to lack of data in many studies on male pubertal milestones. We previously reported a GWAS for recalled age at voice breaking in men (N=55,871) from a single study, 23andMe, which identified 14 genetic independent genetic association signals, and many signals with similar effect sizes on voice breaking as for AAM in women^6^. This overlapping genetic architecture between puberty timing in males (in 23andMe) and females was also reported by others^7^, and supports the use of recalled age at voice breaking in men as an informative measure of puberty timing for further genetic studies.

Although the overall shared genetic architecture for pubertal timing between sexes is high (r_g_=0.74) and many genetic variants show similar effects sizes in both sexes, there are a number of genetic signals that differ between sexes. Most notably, at the *SIM1*/*MCHR2* locus, the allele that promotes earlier puberty in one sex delays it in the other^6^. Sex-specific expression patterns for several of such highlighted genes was reported in the hypothalamus and pituitary of pre-pubertal mice^8^. Further work to identify the pubertal mechanisms that are divergent between sexes may shed light on differences in later life disease risk associations between sexes.

Here, we greatly extend our previously reported GWAS for age at voice breaking in 23andMe^6^ by combining additional data from the UK Biobank study^9^. In this four-fold larger sample, we increase the number of genomic loci that have male-specific effects on puberty timing from 5 to 29, and identify novel biological pathways that warrant further investigation.

## Results

Data on relative age of voice breaking and relative age of first facial hair were available in up to 207,126 male participants in the UK Biobank (UKBB) study. For each of these measures, participants were asked if the event occurred relative to their peers “younger than average”, “about average” or “older than average”. We previously reported that signals for AAM in women show concordant associations with dichotomised voice breaking traits in these UKBB men^5^, but to our knowledge no genetic study has previously evaluated timing of facial hair appearance as a marker of pubertal development.

### Recalled onset of facial hair as a marker of male puberty timing

We defined two dichotomous facial hair variables in UKBB men: 1) relatively-early onset (N=13,226) vs. average onset (N=161,175); and 2) relatively-late onset (N= 26,066) vs. average onset. These facial hair traits were in concordance with similarly dichotomised voice breaking traits defined in the same individuals, although more men reported early or late facial hair onset (39,292; 19.6%) than early or late voice breaking (19,579; 10.2%) **(Supplementary Table 1**). To test the ability of these facial hair traits to detect puberty timing loci, we assessed their associations with previously reported AAM loci. Of the 328 reported autosomal AAM signals for which genotype data were available in UKBB, 274 (83.5%, binomial-P=1.2×10^−33^) and 276 (84.1%, P=7.7×10^−35^) showed directionally-concordant individual associations with relatively-early and relatively-late facial hair, respectively. Furthermore, substantially more AAM signals showed at least nominally significant associations (P<0.05) with relatively-early (102, 31.1%) or relatively-late facial hair (152, 46.3%) compared to ~16 expected by chance for each outcome (**Supplementary Table 2**).

### A multi-trait GWAS for puberty timing in men

We analysed four GWAS models in UKBB men (imputation v2, ~7.4M SNPs): two models (early and late) for each of relative timing of voice breaking and facial hair. The shared genetic architecture between these four traits was high (genetic correlations (r_g_) ranged 0.57 to 0.91; **Supplementary Table 3**), and all four showed high genetic correlation with the continuously measured age at voice breaking in 23andMe (r_g_ 0.61 to 0.81). These high correlations supported the rationale to combine the GWAS data across all five strata using MTAG^10^. This approach enables genetic data from correlated traits and overlapping samples to be combined in a single analysis, which here yielded an effective GWAS meta-analysis sample size of 205,354 men for continuous pubertal timing.

In this combined MTAG dataset, we identified 7,897 variants associated with male puberty timing at genome-wide statistical significance (P<5×10^−8^), comprising 78 independent signals (**Figure 1**; **Supplementary Table 4**). The most significantly associated variant (rs11156429, P=3.5×10^−52^) was located in/near *LIN28B*, consistent with previously reported studies in men and women. Of these 78 signals, 29 were not in linkage disequilibrium (LD, conservatively defined as r^2^<0.05) with a previously reported signal for puberty timing (either AAM in women or voice breaking in men) and were therefore considered novel (**Table 1**).

**Figure 1.**
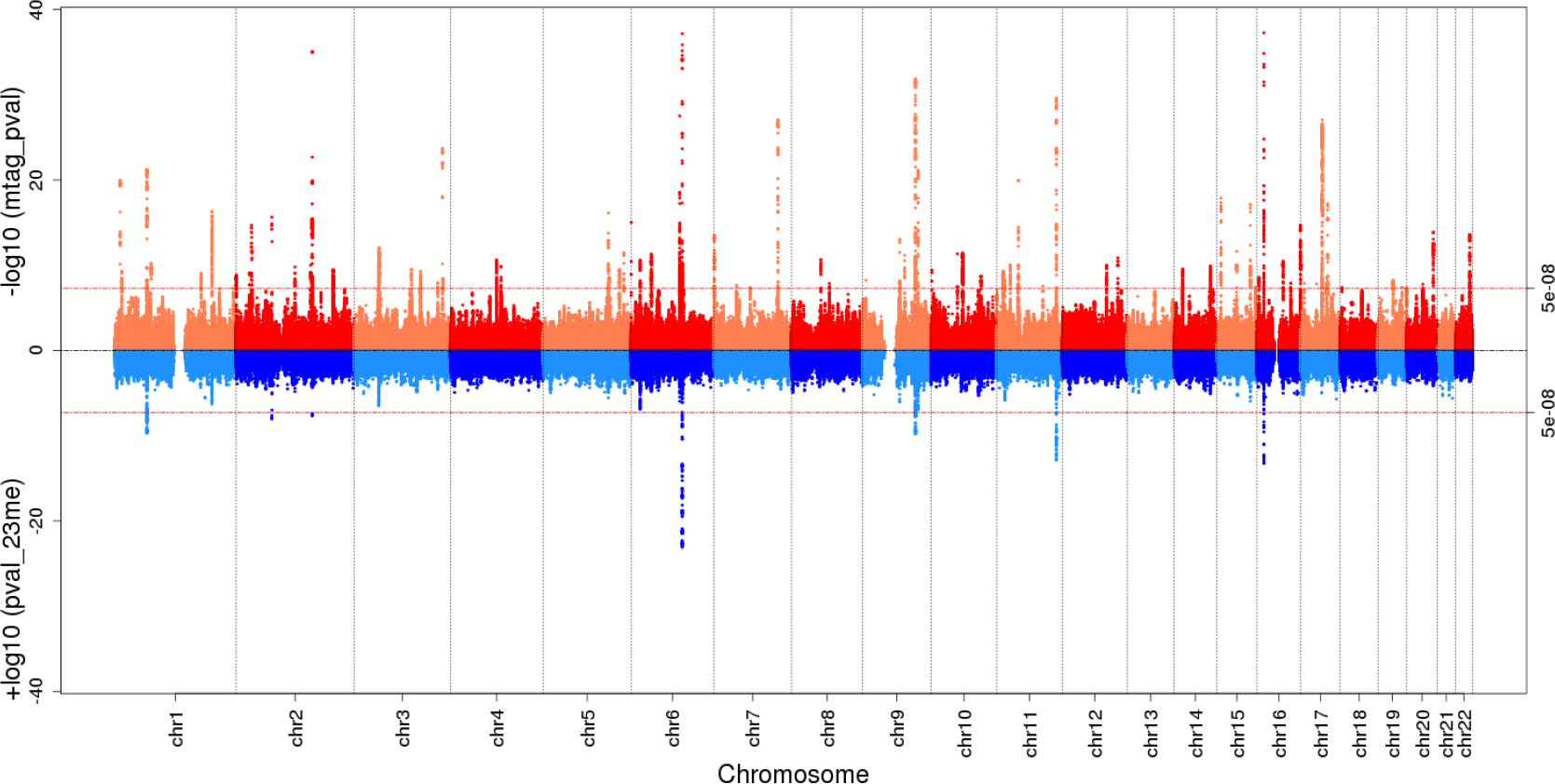
Miami plot displaying multi-trait GWAS (MTAG) −log_10_(P values) for SNP associations with male puberty timing (top half, red shades) and previously reported −log_10_(P values) for age at voice breaking from the 23andMe study GWAS (bottom half, blue shades). Red dashed lines indicate the genome-wide statistical significance threshold (P<5 × 10^−8^).

**Table 1:** Twenty nine novel signals for puberty timing.

| Variant | Chr | Position | Alleles | EAF | Nearest Gene | Voice breaking (MTAG)<sup>1</sup> |  | Menarche (Day *et al.*)<sup>2</sup> |  |
| --- | --- | --- | --- | --- | --- | --- | --- | --- | --- |
|  |  |  |  |  |  | Beta (s.e.) | P | Beta (s.e.) | P |
| rs71578952 | 7 | 131,001,466 | C/T | 0.495 | *MKLN1* | 0.035 (0.003) | $8.4 \times 10^{-28}$ | 0.000 (0.004) | 0.94 |
| rs2222746 | 17 | 44,222,019 | T/G | 0.165 | *KIAA1267* | 0.048 (0.004) | $9.0 \times 10^{-28}$ | n/a | n/a |
| rs73182377 | 3 | 181,512,034 | C/T | 0.227 | *SOX2OT* | 0.040 (0.004) | $1.9 \times 10^{-24}$ | 0.010 (0.005) | 0.05 |
| rs3824915 | 11 | 44,331,509 | G/C | 0.496 | *ALX4* | 0.030 (0.003) | $1.0 \times 10^{-20}$ | 0.008 (0.004) | 0.05 |
| rs77578010 | 1 | 11,035,758 | A/G | 0.776 | *C1orf127* | 0.036 (0.004) | $1.1 \times 10^{-20}$ | 0.006 (0.005) | 0.18 |
| rs7402990 | 15 | 28,384,491 | G/A | 0.916 | *HERC2* | 0.051 (0.006) | $1.4 \times 10^{-18}$ | 0.011 (0.008) | 0.17 |
| rs17833789 | 17 | 55,230,628 | C/A | 0.547 | *AKAP1* | 0.028 (0.003) | $5.9 \times 10^{-18}$ | -0.001 (0.004) | 0.83 |
| rs12203592 | 6 | 396,321 | C/T | 0.170 | *IRF4* | 0.035 (0.004) | $9.6 \times 10^{-16}$ | 0.002 (0.006) | 0.68 |
| rs35063026 | 16 | 89,736,157 | T/C | 0.069 | *C16orf55* | 0.051 (0.006) | $2.1 \times 10^{-15}$ | -0.007 (0.007) | 0.34 |
| rs9690350 | 7 | 547,800 | C/G | 0.419 | *PDGFA* | 0.025 (0.003) | $3.2 \times 10^{-14}$ | -0.011 (0.008) | 0.16 |
| rs6560353 | 9 | 76,375,544 | G/T | 0.161 | *ANXA1* | 0.033 (0.004) | $9.7 \times 10^{-14}$ | 0.016 (0.005) | $2.3 \times 10^{-3}$ |
| rs7905367 | 10 | 54,334,653 | G/C | 0.219 | *MBL2* | 0.027 (0.004) | $4.7 \times 10^{-12}$ | -0.011 (0.005) | 0.03 |
| rs7136086 | 12 | 114,129,719 | C/T | 0.734 | *RBM19* | 0.025 (0.004) | $1.4 \times 10^{-11}$ | 0.019 (0.005) | $3.2 \times 10^{-5}$ |
| rs10110581 | 8 | 60,691,207 | G/A | 0.263 | *CA8* | 0.025 (0.004) | $2.3 \times 10^{-11}$ | 0.014 (0.004) | $1.3 \times 10^{-3}$ |
| rs2842385 | 6 | 19,078,274 | G/A | 0.193 | *MIR548A1* | 0.027 (0.004) | $2.5 \times 10^{-11}$ | 0.021 (0.005) | $1.9 \times 10^{-5}$ |
| rs11836880 | 12 | 91,243,529 | G/C | 0.040 | *C12orf37* | -0.053 (0.008) | $1.1 \times 10^{-10}$ | 0.006 (0.010) | 0.55 |
| rs10164550 | 2 | 121,159,205 | A/G | 0.657 | *INHBB* | 0.022 (0.003) | $1.7 \times 10^{-10}$ | 0.016 (0.004) | $1.1 \times 10^{-4}$ |
| rs12895406 | 14 | 36,998,950 | A/G | 0.536 | *NKX2-1* | 0.021 (0.003) | $2.8 \times 10^{-10}$ | 0.007 (0.004) | 0.05 |
| rs835648 | 3 | 136,671,504 | A/T | 0.696 | *NCK1* | 0.022 (0.004) | $5.2 \times 10^{-10}$ | 0.006 (0.004) | 0.20 |
| rs780094 | 2 | 27,741,237 | T/C | 0.413 | *GCKR* | 0.020 (0.003) | $7.8 \times 10^{-10}$ | 0.018 (0.004) | $3.7 \times 10^{-6}$ |
| rs1979835 | 5 | 135,689,839 | A/G | 0.876 | *TRPC7* | 0.029 (0.005) | $2.5 \times 10^{-9}$ | 0.024 (0.006) | $2.7 \times 10^{-5}$ |
| rs12930815 | 16 | 4,348,635 | C/T | 0.521 | *TFAP4* | 0.019 (0.003) | $2.6 \times 10^{-9}$ | 0.008 (0.004) | 0.03 |
| rs12940636 | 17 | 53,400,110 | C/T | 0.655 | *HLF* | 0.020 (0.003) | $3.6 \times 10^{-9}$ | 0.023 (0.004) | $1.4 \times 10^{-8}$ (3) |
| rs60856990 | 17 | 7,337,853 | A/G | 0.629 | *TMEM102* | 0.019 (0.003) | $1.3 \times 10^{-8}$ | -0.005 (0.004) | 0.19 |
| rs17193410 | 17 | 32,474,149 | G/A | 0.880 | *ACCN1* | 0.028 (0.005) | $1.5 \times 10^{-8}$ | 0.000 (0.007) | 0.98 |
| rs11761054 | 7 | 46,076,649 | C/G | 0.291 | *IGFBP3* | -0.020 (0.004) | $2.2 \times 10^{-8}$ | 0.003 (0.004) | 0.51 |
| rs10765711 | 11 | 94,879,318 | C/G | 0.415 | *ENDOD1* | 0.018 (0.003) | $2.9 \times 10^{-8}$ | -0.001 (0.004) | 0.81 |
| rs61168554 | 15 | 99,286,980 | A/G | 0.361 | *IGF1R* | 0.019 (0.003) | $3.8 \times 10^{-8}$ | 0.017 (0.004) | $2.7 \times 10^{-5}$ |
| rs12983109 | 19 | 49,579,710 | G/A | 0.742 | *KCNA7* | 0.020 (0.004) | $3.8 \times 10^{-8}$ | 0.002 (0.005) | 0.71 |

<sup>1</sup>Current data from multi-trait GWAS for age at voice breaking in men<sup>2</sup>Previously reported GWAS for age at menarche in women (Ref 5)<sup>3</sup>The previous study used a distance based metric to define loci rather than a LD based metric, and only reported robust secondary signals.
Alleles (effect/other); EAF: effect allele frequency; Beta: years per allele; P: P-value from additive models

We sought collective confirmation of the 78 independent signals for male puberty timing in up to 2,394 boys in a prospective birth cohort study, The Avon Longitudinal Study of Parents and Children (ALSPAC)^11^. A polygenic risk score based on the 78 male puberty timing signals was associated with timing of voice breaking in ALSPAC (P_min_=1.6×10^−8^, adjusted r^2^~2%), in addition to other longitudinally assessed pubertal sexual characteristics and growth traits (**Supplementary Table 5**).

### Genetic heterogeneity between sexes

Consistent with our previous study^6^, we observed a moderately strong genome-wide genetic correlation in pubertal timing between males and females (r_g_=0.68, P=2.6×10^−213^; based on continuous data on voice breaking and AAM in 23andMe), with similar effect estimates in both sexes for many individual variants (**Figure 2**). However, there were exceptions to this overall trend: 5/78 male puberty timing signals (**Figure 2b**) and 15/387 reported AAM signals (**Figure 2a**) showed significant (by Bonferroni corrected P-values) heterogeneity between sexes in their effects on puberty timing (two of these heterogeneous signals were found in both analyses) **(Supplementary Tables 6 and 7**; **Figure 2**). Only one signal showed significant directionally-opposite effects (i.e. the allele that conferred earlier puberty timing in one sex delayed puberty in the other sex); rs6931884 at *SIM1/PRDM13/MCHR2*, as previously reported (males: **β**_voice-breaking_= −0.064 years/allele; females: **β**_menarche_=0.059 years/allele; P_heterogeneity_=2.6×10^−14^). Two variants located near to genes that are disrupted in rare disorders of puberty showed no effect or weaker effect in males than in females: rs184950120, 5’UTR to *MKRN3* (**β**_voice-breaking_=0.085 years/allele; **β**_menarche_=0.396 years/allele, P_heterogeneity_=3.6×10^−3^), and rs62342064, one of 3 AAM variants in/near *TACR3* (**β**_voice-breaking_= −0.017 years/allele; **β**_menarche_=0.057 years/allele, P_heterogeneity_=4.2×10^−5^).

**Figure 2.**
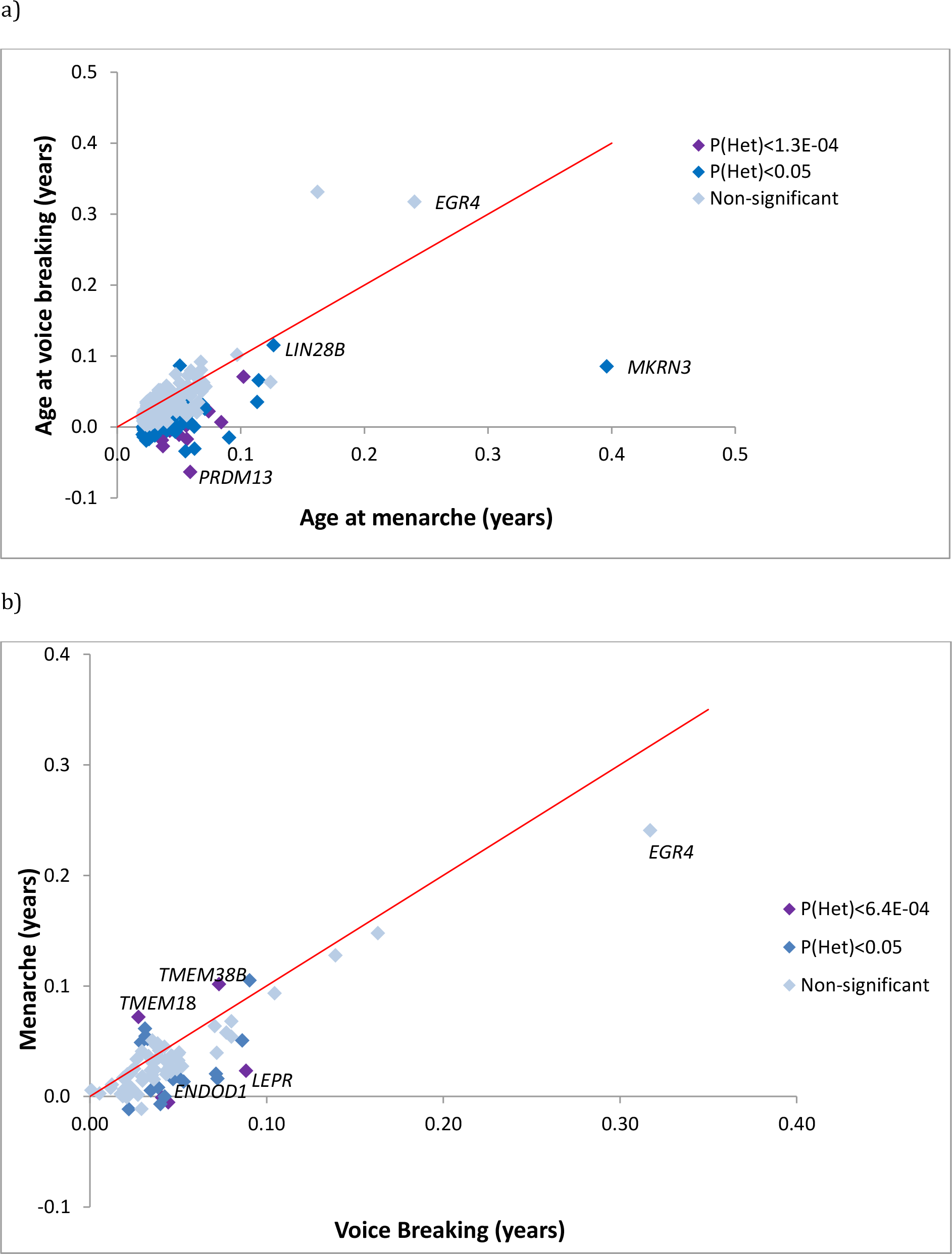
Scatterplots comparing effect sizes on age at menarche (in women) and age at voice breaking (in men) for: **a)** known menarche loci and **b)** male puberty loci identified in the current meta-analysis. SNPs are not independent across the two panels. Variants are coloured based on heterogeneity (P-value) between women and men. Red lines indicate the perfect agreement (X=Y).

### Implicated genes, tissues and biological pathways

Two of the 78 lead variants associated with male pubertal timing were non-synonymous: a previously reported AAM signal in *KDM4C* (rs913588), encoding a lysine-specific demethylase, and a novel male-specific signal in *ALX4* (rs3824915), encoding a homeobox gene involved in fibroblast growth factor (FGF) signalling that is mutated in rare disorders of cranium/central neural system (CNS) development with male-specific hypogonadism^12^. A further 10 lead variants were in strong LD (r^2^>0.8) with one or more non-synonymous variants, of which three represent novel signals for puberty timing: *FGF11*, which encodes a FGF expressed in the developing CNS and promotes peripheral androgen receptor expression^13^, *TFAP4*, which encodes a transcription factor of the basic helix-loop-helix-zipper family^14^, and *GCKR*, which encodes a regulatory protein that inhibits glucokinase in liver and pancreatic islets and is associated with a range of cardio-metabolic traits15 (**Supplementary Table 8**). A further 7 are reported signals for AAM, but were not previously reported for voice-breaking. These missense variants are in the following genes: *SRD5A2*, encoding for Steroid 5-alpha-reductase, which converts testosterone into the more potent androgen dihydrotestosterone; *LEPR*, encoding the receptor for appetite and reproduction hormone leptin; *SMARCAD1*, encoding a mediator of histone H3/H4 deacetylation; *BDNF, FNDC9, FAM118A*, and *ZNF446*.

Consistent with genetic analyses of AAM in females, tissues in the CNS were the most strongly enriched for genes co-located near to male puberty timing associated variants (**Supplementary Figures 1 and 2; Supplementary Tables 9 and 10**). To identify mechanisms that regulate pubertal timing in males, we tested all SNPs genome-wide for enrichment of voice breaking associations with pre-defined biological pathway genes. Four pathways showed evidence of enrichment: histone methyltransferase complex (FDR=0.01); regulation of transcription (FDR=0.02)); ATP binding (FDR=0.03); and cAMP biosynthetic process (FDR=0.03) (**Supplementary Table 11**).

### Genetic and phenotypic links between hair colour and puberty timing

Noting that three novel loci for puberty timing were located proximal to genes previously associated with pigmentation (*HERC2*, *IRF4*, *C16orf55*), we assessed the broader relationship between these traits. It is known that men have darker natural hair colour than women in European ancestry populations^16^ and this sex difference appears following the progressive darkening of hair and skin colour during adolescence^17,18^. However, a link between inter-individual variation in natural hair colour and puberty timing has not previously been described. We assessed this phenotypic relationship in up to 179,594 white males of European ancestry in UKBB in a model including 40 genetic principal components (to adjust for even minor subpopulation ethnic variations). Men with red, dark brown and black natural hair colours showed progressively higher odds of early puberty timing, relative to men with blond hair. Similarly, women with darker natural hair colours had earlier puberty timing relative to women with blond hair (**Table 2**).

**Table 2:** Phenotypic associations between natural hair colour and puberty timing in UK Biobank men and women.

| Natural hair colour | Men (n=179,549) |  |  | Women (n=238,195) |  |  |
| --- | --- | --- | --- | --- | --- | --- |
|  | Effect* | 95% CI | P-value | Effect* | 95% CI | P-value |
| Blond | (ref) |  |  | (ref) |  |  |
| Red | 1.17 | 1.02, 1.35 | 0.02 | -0.100 | -0.134, -0.066 | $6.2 \times 10^{-9}$ |
| Light brown | 1.00 | 0.91, 1.09 | 0.92 | -0.026 | -0.047, -0.005 | 0.01 |
| Dark brown | 1.45 | 1.34, 1.58 | $6.7 \times 10^{-18}$ | -0.093 | -0.114, -0.072 | $6.7 \times 10^{-18}$ |
| Black | 1.63 | 1.46, 1.81 | $1.2 \times 10^{-19}$ | -0.059 | -0.113, -0.005 | 0.03 |

\*Effect size in men is odds ratio for early relatively voice breaking (relative to blond hair). Effect size in women is mean difference in age at menarche in years (relative to blond hair).

To explore the genetic links between these two phenotypes, we systematically assessed the effects of genetic variants associated with natural hair colour on puberty timing. Using a two-sample Mendelian randomization (MR) approach, we modelled 119 recently reported GWAS hair colour signals as a single instrumental variable16 on age at voice breaking (in 23andMe men). The findings inferred that darker natural hair colour confers earlier puberty timing in men (**β**_IVW_= −0.044 years per ordered category, P_IVW_=7.10×10^−3^) with no evidence of heterogeneity across signals (Cochrane Q: P=0.99) or directional pleiotropy (MR-Egger Intercept: P=0.99) (**Supplementary Figure 3**). Using a similar approach in published AAM data in females5, we inferred a directionally consistent but weaker effect of darker natural hair colour on earlier AAM (**β**_IVW_= −0.017, PIVW=3.64×10^−3^), and with modest heterogeneity across signals (Cochrane Q: P=0.038) (**Supplementary Table 12**).

Due to the partial overlapping samples between the SNP instrument discovery and outcome in the above approach, we performed a second, more conservative analysis, using a more limited 5-SNP score aligned to darker hair colour in non-overlapping data on white UK Biobank individuals, adjusting for 40 genetic principal components and geographical location of testing centres. This found highly consistent results in men: the 5-SNP score was associated with higher risk for early voice breaking (P=1.72×10^−19^) and lower risk for late voice breaking (P=5.90×10^−6^); but we found no association with AAM in women (P=0.23) (**Supplementary Table 12**).

### Relevance of male puberty timing to other complex traits and diseases

To assess the extent of shared heritability between male puberty timing and other complex traits, we calculated genome-wide genetic correlations across 751 complex traits/datasets using LD score regression^19^ (**Supplementary Table 13**). Apart from other puberty and growth-related traits, the strongest positive genetic correlation was observed with ‘overall health rating’, followed by several social traits, including educational attainment, fluid intelligence score and ages at first/last birth. Conversely, male puberty timing showed negative genetic correlations with cardiometabolic diseases, including Type 2 diabetes and hypertension, as well as health risk behaviours, including alcohol intake frequency and smoking **(Supplementary Figure 4**). In general, early genetically-predicted puberty in men was correlated with adverse health outcomes. We then performed MR analyses to test the causal relationships between male puberty timing on two exemplar traits: lifespan and prostate cancer.

We previously reported causal relationships between earlier AAM and higher risks for hormone sensitive cancers, when adjusting for the protective effects of higher BMI^5,20^. Here, using a similar two-sample multi-variate MR approach, we found that adjustment for BMI revealed apparent protective effects of later male puberty on the risk of overall prostate cancer (OR per year=0.89, 95%CI=0.79-1.00; P=0.043) or advanced prostate cancer (OR per year=0.82, 95%CI=0.67-1.00; P=0.048) (**Supplementary Table 14; Supplementary Figure 5)**. We used a similar causal modelling approach to test the relevance of male puberty timing to lifespan (see Methods). The findings support a causal effect of later puberty timing in males on longer lifespan, corresponding to 7 months longer life per year later puberty (P=7.0×10^−5^) (**Figure 3**).

**Figure 3:**
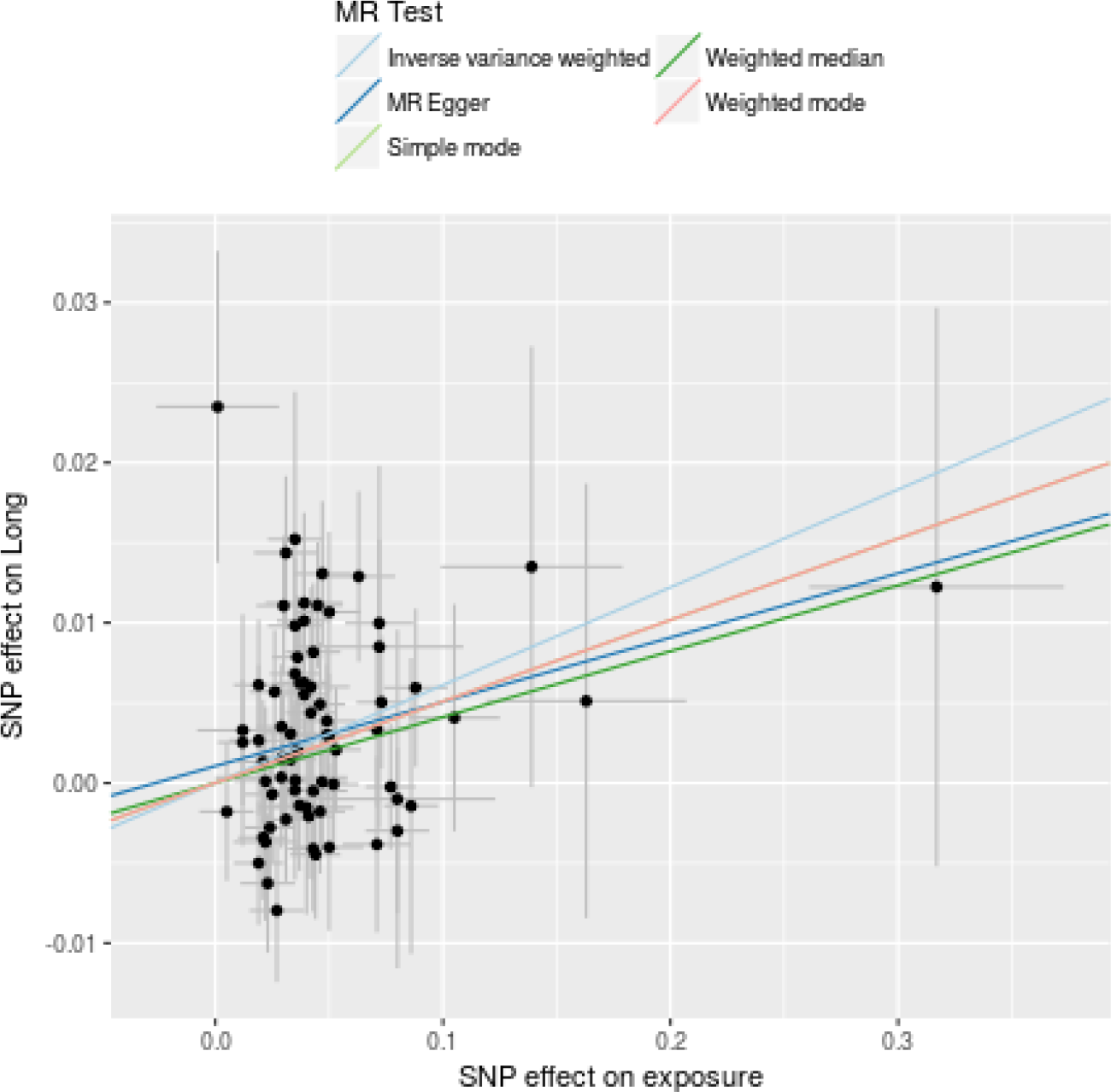
Scatterplot of 75 male puberty loci (pruned for heterogeneity) comparing effect sizes for puberty timing and longevity. Lines show results for different MR models. MR-Egger test showed no evidence for directional pleiotropy (Egger Intercept=0.001, P=0.44).

## Discussion

Here, we report the largest genetic study to date for male puberty timing by performing a meta-analysis across closely related phenotypic traits in two large studies. The effective sample size of 205,354 men is substantially higher than the previous GWAS in men (n=55,871), although still smaller than similar studies in women (n~370,000). While there was overall moderately strong overlap between sexes in the genetic architecture of puberty timing (r_g_=0.68), several findings point to mechanisms with particular relevance to men. Newly highlighted genes implicated in puberty timing include, *ALX4* and *SRD5A2*, the disruption of which leads to male-specific reproductive disorders, and *INHBB* encoding the beta-B subunit of the hormone Inhibin B, which is secreted by testicular Sertoli cells in males. By contrast, the known female puberty timing locus at *INHBA*, which encodes the beta-A subunit of the hormone Inhibin A, showed little association with puberty timing in males.

Non-genetic observational studies have consistently reported darkening of hair and skin pigmentation in children of European ancestry during the peri-pubertal years^17,18^. Furthermore, the established sexual dimorphism in pigmentation^16^ reportedly appears from puberty onwards^17^ and the relatively darker skin and hair of men compared to women is postulated to reflect the stronger stimulation of melanogenesis by androgens compared to estrogens^21^. Our findings suggest a more widespread overlap between pigmentation and reproduction, possibly reflecting common regulators of pituitary production of melanocortins and gonadotrophins, or even an impact of melanocyte signalling on puberty timing. The pituitary pro-peptide, pro-opiomelanocortin (POMC), is cleaved into several peptides with significant melanogenic activity (ACTH, α-MSH, and β-MSH) by the pro-hormone convertase enzymes, PC-1 and PC-2, both of which are implicated by previously reported loci for AAM in females^5^. The relationship in women is more complicated suggesting that while there is a relationship between pigmentation and puberty timing in both sexes, it may act in a sex-specific manner.

Finally, our findings substantiate links between puberty timing in men and later life health outcomes, including overall life span. Specifically, as recently described for risks of breast, ovary and endometrial cancers in women^5^, here we inferred a causal effect of earlier puberty timing on higher risk of prostate cancer in males, indicative of some BMI-independent mechanism(s), such as higher sex steroid exposure. In conclusion, our findings demonstrate the utility of multi-trait GWAS to combine data across studies with related measures to provide novel insights into the regulation and consequences of puberty timing.

## Methods

### 23andMe study

Genome-wide SNP data were available in up to 55,871 men aged 18 or older of European ancestry from the 23andMe study. Age at voice breaking was determined by response to the question ‘How old were you when your voice began to crack/deepen?’ in an online questionnaire. Participants chose from one of seven pre-defined age bins (under 9, 9-10 years old, 11-12 years old, 13-14 years old, 15-16 years old, 17-18 years old, 19 years old or older). These were then re-scaled to one year age bins post-analysis by a previously validated method22. SNPs were excluded prior to imputation based on the following criteria: Hardy-Weinberg equilibrium P < 10^−20^; call rate < 95%; or a large frequency discrepancy when compared with European 1000 Genomes reference data6. Imputation of genotypes was performed using the March 2012 ‘v3’ release of 1000 Genomes reference haplotype panel. Genetic associations with puberty timing were obtained by linear regression models using age and five genetically determined principal components to account for population structure as covariates, with additive allelic effects assumed. P values for SNP associations were computed using likelihood ratio tests. Participants provided informed consent to take part in this research under a protocol approved by Ethical and Independent Review Services, an institutional review board accredited by the Association for the Accreditation of Human Research Protection Programs.

### UK Biobank

Genotyping for UK Biobank participants has been described in detail previously^9^. For this analysis, we limited our sample to individuals of white European ancestry. Age at voice breaking for male participants from the UK Biobank cohort study was obtained by responses to the touch-screen question ‘When did your voice break?’ Participants were required to choose from one of five possible options (younger than average, about average, older than average, do not know, prefer not to answer). For age at first facial hair, respondents were asked to choose from one of the same five options in response to the touch-screen question ‘When did you start to grow facial hair?’ In total, data were available in up to 191,270 individuals for voice breaking and 198,731 individuals for facial hair.

Respondents who answered either ‘older than average’ or ‘younger than average’ were compared separately using the ‘about average’ group as a reference in a case-control design for both phenotypes. For both voice breaking and facial hair, relatively-early and relatively-late effect estimates were obtained using linear mixed models which were applied using BOLT-LMM software, which accounts for cryptic population structure and relatedness. Covariates included age, genotyping chip and 10 principle components.

### Meta-analysis of voice breaking results

GWAS summary results for each of the five strata (23andMe age at voice breaking; UK Biobank relatively-early and late voice breaking; and UK Biobank relatively-early and late facial hair) were meta-analysed using MTAG^10^. MTAG uses GWAS summary statistics from multiple correlated traits to effectively increase sample size and statistical power to detect genetic associations. Full details on the methodology have been described previously^10^. In brief, MTAG estimates a variance-covariance matrix to correlate the effect sizes of each trait using a maximum likelihood method, with each trait and genotype standardised to have a mean of zero and variance of one. In addition, MTAG calculates a variance-covariance matrix for the GWAS estimation error using LD score regressions. The effect estimate for the association of each SNP on the trait of interest is then derived using a marginal likelihood function, in a generalisation of standard inverse-variance weighted meta-analysis.

Prior to meta-analysis, we removed extremely rare variants (MAF<0.01). In addition, for the four UK Biobank strata we calculated the effective sample size using the following equation:

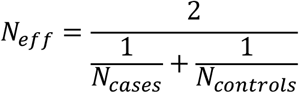

Effective sample sizes for early and late voice breaking were 15,711 and 21,217, respectively and 17,391 and 23,011 for early and late facial hair, respectively. We used a conservative genome-significant P-value threshold of P<5 × 10^−8^ to determine significant SNP associations. Independent signals were identified using distance-based clumping, with the SNP with the lowest P-value within a 1 MB window being considered the association signal at that locus.

### Gene annotation and identification of novel loci

For each independent signal identified in the voice breaking meta-analysis, we identified all previously reported age at menarche (AAM) and voice breaking (VB) loci within 1MB of that SNP. A locus was considered novel if there were no previously reported puberty loci (for AAM or VB) within 1MB, or if no loci within 1MB were in LD with it (*r*^2^<0.05). Heterogeneity between AAM and VB for each SNP was determined by the I2 statistic and P-value generated by METAL software.

Gene annotation was performed using a combination of methods. Information on the nearest annotated gene was obtained from HaploReg v4.1. In addition, other genes in the region were identified using plots produced from LocusZoom. The most likely causal variant was determined by combining this information with identification of any non-synonymous variants within the region as well as application of existing knowledge.

### Replication in ALSPAC

The Avon Longitudinal Study of Parents and Children (ALSPAC) recruited pregnant women resident in the Avon area of the UK with an expected delivery date between 1^st^ April 1991 and 31^st^ December 1992. Since then, mothers, partners, and offspring have been followed up regularly through questionnaires and clinical assessments^11^. The offspring cohort consists of 14,775 live-born children (75.7% of the eligible live births). Full details of recruitment, follow-up and data collection have been reported previously^11^. Ethical approval for the study was obtained from the ALSPAC Ethics and Law committee and the Local Research Ethics Committees. A series of nine postal questionnaires regarding pubertal development was administered approximately annually from the time the participant was aged 8 until he was aged 17. The questionnaires, which were responded by either the parents or the participant, had schematic drawings and verbal descriptions of secondary sexual characteristics (genitalia and pubic hair development) based on the Tanner staging system, as well as information on armpit hair growth and voice change. Age at voice change was considered the age at which the adolescent reported his voice to be occasionally a lot lower or to have changed completely. Weight and height were measured annually up to age 13 years, then at ages 15 and 17 years by a trained research team. Age at peak height velocity (PHV) was estimated using Superimposition by Translation And Rotation (SITAR) mixed effects growth curve analysis^23^. The sample size available varied according to the phenotype, from 1,126 (genital development at which armpit hair started to grow).

The genetic risk score (GRS) was calculated based on 73 SNPs weighted by the effect size reported for that SNP in the ReproGen Consortium. The GRS was standardized, and results are presented as increase in the phenotype per standard-deviation increase in the GRS. Linear (continuous phenotype) and logistic (binary phenotype) regression analyses were performed unadjusted and adjusted for age (except for age at PHV, age at voice change and age at which armpit hair started to grow) and controlled for population stratification using the first 10 principal components.

### Gene expression and pathway analysis

We used MAGENTA to investigate whether genetic associations in the meta-analysed dataset showed enrichment in any known biological pathways. MAGENTA has previously been described in detail^24^. In brief, genes are mapped to an index SNP based on a 150 kb window, with a regression model applied to correct the P-value (gene score) for gene size, SNP density and LD-related properties. Gene scores are ranked, and the numbers of gene scores observed in a given pathway in the 75^th^ and 95^th^ percentiles are calculated. A P-value for gene-set enrichment analysis (GSEA) is calculated by comparing these values to one million randomly generated gene sets. Testing was completed on 3,216 pathways from four databases (PANTHER, KEGG, Gene Ontology and Ingenuity). Significance was determined based on an FDR<0.05 for genes in the 75^th^ or 95^th^ percentile.

To determine tissue-specific expression of genes, we used information from the GTEx project. GTEx characterizes transcription levels of RNA in a variety of tissue and cell types, using sample from over 1,000 deceased individuals of European, African-American and Asian descent. We investigated transcription levels of significant genes identified in our meta-analysis of voice breaking in 53 different tissue types. We used a conservative Bonferroni-corrected P-value of 9.4×10^−4^ (=0.05/53) to determine significance.

### Association between hair colour and puberty timing

Information on natural hair colour for UK Biobank participants was collected via touchscreen questionnaire, in response to the question “What best describes your natural hair colour? (If your hair colour is grey, the colour before you went grey)”. Participants chose from one of 6 possible colours: blond, red, light brown, dark brown, black or other. For our analyses, we restricted this to include only non-related individuals of white European ancestry, totalling 190,845 men and 238,179 women. Hair colours were assigned numerical values from lightest (blond) to darkest (black) in order to perform ordered logistic regression of hair colour for both relative age at voice breaking in men and AAM in women. In both cases, blond hair was used as the reference group and models were adjusted for the top 10 principle components to account for population structure. In men this produces an effect estimate as an odds ratio for early puberty (relative to blond-haired individuals), while in women the effect estimate is on a continuous scale for AAM (in years) relative to the mean AAM for those with blond hair.

### Genetic correlations

Genetic correlations (r_g_) were calculated between age of puberty in males and 751 health-related traits which were publically available from the LD Hub database using LD Score Regression^19,25^.

### Mendelian randomisation analyses

#### Longevity

GWAS summary statistics for longevity were obtained from Timmers et al. (paper under consideration). Briefly, Timmers et al. performed a GWAS of parent survivorship under the Cox proportional hazard model in 1,012,050 parent lifespans of unrelated subjects using methods of Joshi et al., but extending UK Biobank data to that of second release. Data were meta-analysed using inverse variance meta-analysis results from UK Biobank genomically British, Lifegen, UK Biobank self-reported British (but not identified as genomically British), UK Biobank Irish, and UK Biobank other white European descent. The resultant hazard ratios and their standard errors were then taken forward for two-sample MR using the 78 male puberty loci and MR-Base.

#### Prostate cancer

Summary statistics for the association between the genetic variants and risk of prostate cancer were obtained from the PRACTICAL/ELLIPSE consortium, based on GWAS analyses of 65,044 prostate cancer cases and 48,344 controls (all of European ancestry) genotyped using the iCOGS or OncoArray chips^26^. The analyses were repeated using summary statistics from a comparison of the subset of 9,640 cases with advanced disease versus 45,704 controls, where advanced cases were defined as those with at least one of: Gleason score 8+, prostate cancer death, metastatic disease or PSA>100. Two-sample MR analyses were conducted using weighted linear regressions of the SNP-prostate cancer log odds ratios (logOR) on the SNP-puberty beta coefficients, using the variance of the logORs as weights. This is equivalent to an inverse-variance weighted meta-analysis of the variant-specific causal estimates. Because of evidence of over-dispersion (i.e. heterogeneity in the variant-specific causal estimates), the residual standard error was estimated, making this equivalent to a random-effects meta-analysis. Unbalanced horizontal pleiotropy was tested based on the significance of the intercept term in MR-Egger regression. The total effect of puberty timing on prostate cancer risk was separated into a direct effect (independent of BMI, a potential mediator) and an indirect effect (operating via BMI), as described in Burgess *et al.*^19^

#### Hair colour

Causal associations between hair colour and puberty timing were assessed in two ways. First, we performed a two-sample MR analysis based on summary statistics (as described in Burgess *et al.*^19^) using the most recently reported GWAS for hair colour^16^ and 23andMe data for SNP effect estimates on age at voice breaking in men, and published ReproGen consortium data on AAM in women. However, there is partial overlap (between samples used for the discovery phase GWAS of hair colour variants (n=290,891 from 23andMe plus UK Biobank) and the samples used for puberty timing (n=55,871 men from 23andMe; maximum potential overlap=55,871/290,891=19%). Therefore, we also performed a sensitivity analysis in non-overlapping samples; we used a more limited 5-SNP instrument for darker hair colour (identified in an earlier hair colour GWAS that did not include UK Biobank^27^), and assessed its effects on puberty timing in UK Biobank men and women in an individual-level MR analysis, controlling for geographical (assessment centre) and genetic ancestry factors (40 principal components).

## Supporting information

## Acknowledgements

This work was funded by the UK Medical Research Council (MRC - Unit programme MC_UU_12015/2 and used UK Biobank data under application 9905. The MRC and Wellcome (Grant ref: 102215/2/13/2) and the University of Bristol provide core support for ALSPAC. This publication is the work of the authors and <John Perry and Ken Ong> will serve as guarantors for the contents of this paper. GWAS data for ALSPAC was generated by Sample Logistics and Genotyping Facilities at Wellcome Sanger Institute and LabCorp (Laboratory Corporation of America) using support from 23andMe. We are extremely grateful to all the families who took part in this (ALSPAC) study, the midwives for their help in recruiting them, and the whole ALSPAC team, which includes interviewers, computer and laboratory technicians, clerical workers, research assistants, volunteers, managers, receptionists and nurses. We thank the research participants and employees of 23andMe for making this work possible.

## Data Availability

### 23andMe

Full summary statistics for 23andMe datasets will be made available to qualified researchers under an agreement that protects participant privacy. Researchers should visit https://research.23andme.com/dataset-access/ for more details and instructions for applying for access to the data.

### ALSPAC

Please note that the study website contains details of all the data that is available through a fully searchable data dictionary and variable search tool at http://www.bristol.ac.uk/alspac/researchers/our-data/.

