## Supplementary material for "Expanded genomic analyses for male voice-breaking highlights a shared phenotypic and genetic basis between puberty timing and hair colour"

**ADDITIONAL AUTHOR INFORMATION**

Michelle Agee, Babak Alipanahi, Adam Auton, Robert K. Bell, Katarzyna Bryc, Sarah L. Elson, Pierre Fontanillas, Nicholas A. Furlotte, David A. Hinds, Karen E. Huber, Aaron Kleinman, Nadia K. Litterman, Matthew H. McIntyre, Joanna L. Mountain, Elizabeth S. Noblin, Carrie A.M. Northover, Steven J. Pitts, J. Fah Sathirapongsasuti, Olga V. Sazonova, Janie F. Shelton, Suyash Shringarpure, Chao Tian, Joyce Y. Tung, Vladimir Vacic, Catherine H. Wilson.

**The PRACTICAL Consortium (**[**http://practical.icr.ac.uk/**](http://practical.icr.ac.uk/)**):**

Rosalind A. Eeles^1,2^, Brian E. Henderson^3*^, Christopher A. Haiman^3^, ZSofia Kote-Jarai^1^, Fredrick R. Schumacher^4,5^, Ali Amin Al Olama^6,7^, Sara Benlloch^6,1^, Kenneth Muir^8,9^, Sonja I. Berndt^10^, David V. Conti^3^, Fredrik Wiklund^11^, Stephen Chanock^10^, Susan Gapstur^12^, Victoria L. Stevens^12^, Catherine M. Tangen^13^, Jyotsna Batra^14,15^, Judith Clements^14,15^, Australian Prostate Cancer BioResource (APCB)^14^, Henrik Gronberg^11^, Nora Pashayan^16,17^, Johanna Schleutker^18,19^, Demetrius Albanes^10^, Alicja Wolk^20, 21^, Catharine West^22^, Lorelei Mucci^23^, Géraldine Cancel-Tassin^24,25^, Stella Koutros^10^, Karina Dalsgaard Sorensen^26,27^, Eli Marie Grindedal^28^, David E. Neal^29,30,31^, Freddie C. Hamdy^31^, Jenny L. Donovan^32^, Ruth C. Travis^33^, Robert J. Hamilton^34^, Sue Ann Ingles^3^, Barry S. Rosenstein^35,36^, Yong-Jie Lu^37^, Graham G. Giles^38,39^, Adam S. Kibel^40^, Ana Vega^41^, Manolis Kogevinas^42,43,44,45^, Kathryn L. Penney^46^, Jong Y. Park^47^, Janet L. Stanford^48,49^, Cezary Cybulski^50^, Børge G. Nordestgaard^51,52^, Hermann Brenner^53,54,55^, Christiane Maier^56^, Jeri Kim^57^, Esther M. John^58,59^, Manuel R. Teixeira^60,61^, Susan L. Neuhausen^62^, Kim De Ruyck^63^, Azad Razack^64^, Lisa F. Newcomb^48,65^, Davor Lessel^66^, Radka Kaneva^67^, Nawaid Usmani^68,69^, Frank Claessens^70^, Paul A. Townsend^71^, Manuela Gago-Dominguez^72,73^, Monique J. Roobol^74^, Florence Menegaux^75^, Kay-Tee Khaw^76^, Lisa Cannon-Albright^77,78^, Hardev Pandha^79^, Stephen N. Thibodeau^80^

^*^In memorium

^1^ The Institute of Cancer Research, London, UK.

^2^ Royal Marsden NHS Foundation Trust, London, UK.

^3^ Department of Preventive Medicine, Keck School of Medicine, University of Southern California/Norris Comprehensive Cancer Center, Los Angeles, CA, USA.

^4^ Department of Epidemiology and Biostatistics, Case Western Reserve University, Cleveland, OH, USA.

^5^ Seidman Cancer Center, University Hospitals, Cleveland, OH, USA.

^6^ Centre for Cancer Genetic Epidemiology, Department of Public Health and Primary Care, University of Cambridge, Strangeways Research Laboratory, Cambridge, UK.

^7^ University of Cambridge, Department of Clinical Neurosciences, Cambridge, UK.

^8^ Division of Population Health, Health Services Research and Primary Care, University of Manchester, Manchester, UK.

^9^ Warwick Medical School, University of Warwick, Coventry, UK.

^10^ Division of Cancer Epidemiology and Genetics, National Cancer Institute, NIH, Bethesda, MD, USA.

^11^ Department of Medical Epidemiology and Biostatistics, Karolinska Institute, Stockholm, Sweden.

^12^ Epidemiology Research Program, American Cancer Society, 250 Williams Street, Atlanta, GA, USA.

^13^ SWOG Statistical Center, Fred Hutchinson Cancer Research Center, Seattle, WA, USA.

^14^ Australian Prostate Cancer Research Centre-Qld, Institute of Health and Biomedical Innovation and School of Biomedical Science, Queensland University of Technology, Brisbane, Queensland, Australia.

^15^ Translational Research Institute, Brisbane, Queensland, Australia.

^16^ University College London, Department of Applied Health Research, London, UK.

^17^ Centre for Cancer Genetic Epidemiology, Department of Oncology, University of Cambridge, Strangeways Laboratory, Cambridge, UK.

^18^ Department of Medical Biochemistry and Genetics, Institute of Biomedicine, University of Turku, Finland.

^19^ Tyks Microbiology and Genetics, Department of Medical Genetics, Turku University Hospital, Finland.

^20^ Division of Nutritional Epidemiology, Institute of Environmental Medicine, Karolinska Institutet, Sweden.

^21^ Department of Surgical Sciences, Uppsala University, Uppsala, Sweden.

^22^ Division of Cancer Sciences, University of Manchester, Manchester Academic Health Science Centre, Radiotherapy Related Research, Manchester NIHR Biomedical Research Centre, The Christie Hospital NHS Foundation Trust, Manchester, UK.

^23^ Department of Epidemiology, Harvard T.H Chan School of Public Health, Boston, MA, USA.

^24^ CeRePP, Tenon Hospital, Paris, France.

^25^ UPMC Sorbonne Universites, GRC N°5 ONCOTYPE-URO, Tenon Hospital, Paris, France.

^26^ Department of Molecular Medicine, Aarhus University Hospital, Denmark.

^27^ Department of Clinical Medicine, Aarhus University, Denmark.

^28^ Department of Medical Genetics, Oslo University Hospital, Norway.

^29^ University of Cambridge, Department of Oncology, Addenbrooke's Hospital, Cambridge, UK.

^30^ Cancer Research UK Cambridge Research Institute, Li Ka Shing Centre, Cambridge, UK.

^31^ Nuffield Department of Surgical Sciences, University of Oxford, Oxford, UK, Faculty of Medical Science, University of Oxford, John Radcliffe Hospital, Oxford, UK.

^32^ School of Social and Community Medicine, University of Bristol, Bristol, UK.

^33^ Cancer Epidemiology Unit, Nuffield Department of Population Health University of Oxford, Oxford, UK.

^34^ Dept. of Surgical Oncology, Princess Margaret Cancer Centre, Toronto, Canada.

^35^ Department of Radiation Oncology, Icahn School of Medicine at Mount Sinai, New York, NY, USA.

^36^ Department of Genetics and Genomic Sciences, Icahn School of Medicine at Mount Sinai, New York, NY, USA.

^37^ Centre for Molecular Oncology, Barts Cancer Institute, Queen Mary University of London, John Vane Science Centre, London, UK.

^38^ Cancer Epidemiology & Intelligence Division, The Cancer Council Victoria, Melbourne, Victoria, Australia.

^39^ Centre for Epidemiology and Biostatistics, Melbourne School of Population and Global Health, The University of Melbourne, Melbourne, Australia.

^40^ Division of Urologic Surgery, Brigham and Womens Hospital, Boston, MA, USA.

^41^ Fundación Pública Galega de Medicina Xenómica-SERGAS, Grupo de Medicina Xenómica, CIBERER, IDIS, Santiago de Compostela, Spain.

^42^ Centre for Research in Environmental Epidemiology (CREAL), Barcelona Institute for Global Health (ISGlobal), Barcelona, Spain.

^43^ CIBER Epidemiología y Salud Pública (CIBERESP), Madrid, Spain.

^44^ IMIM (Hospital del Mar Research Institute), Barcelona, Spain.

^45^ Universitat Pompeu Fabra (UPF), Barcelona, Spain.

^46^ Channing Division of Network Medicine, Department of Medicine, Brigham and Women's Hospital/Harvard Medical School, Boston, MA, USA.

^47^ Department of Cancer Epidemiology, Moffitt Cancer Center, Tampa, USA.

^48^ Division of Public Health Sciences, Fred Hutchinson Cancer Research Center, Seattle, Washington, USA.

^49^ Department of Epidemiology, School of Public Health, University of Washington, Seattle, Washington, USA.

^50^ International Hereditary Cancer Center, Department of Genetics and Pathology, Pomeranian Medical University, Szczecin, Poland.

^51^ Faculty of Health and Medical Sciences, University of Copenhagen, Denmark.

^52^ Department of Clinical Biochemistry, Herlev and Gentofte Hospital, Copenhagen University Hospital, Herlev, Denmark.

^53^ Division of Clinical Epidemiology and Aging Research, German Cancer Research Center (DKFZ), Heidelberg, Germany.

^54^ German Cancer Consortium (DKTK), German Cancer Research Center (DKFZ), Heidelberg, Germany.

^55^ Division of Preventive Oncology, German Cancer Research Center (DKFZ) and National Center for Tumor Diseases (NCT), Heidelberg, Germany.

^56^ Institute for Human Genetics, University Hospital Ulm, Ulm, Germany.

^57^ The University of Texas MD Anderson Cancer Center, Department of Genitourinary Medical Oncology, Houston, TX, USA.

^58^ Cancer Prevention Institute of California, Fremont, CA, USA.

^59^ Department of Health Research & Policy (Epidemiology) and Stanford Cancer Institute, Stanford University School of Medicine, Stanford, CA , USA.

^60^ Department of Genetics, Portuguese Oncology Institute of Porto, Porto, Portugal.

^61^ Biomedical Sciences Institute (ICBAS), University of Porto, Porto, Portugal.

^62^ Department of Population Sciences, Beckman Research Institute of the City of Hope, Duarte, CA, USA.

^63^ Ghent University, Faculty of Medicine and Health Sciences, Basic Medical Sciences, Gent, Belgium.

^64^ Department of Surgery, Faculty of Medicine, University of Malaya, Kuala Lumpur, Malaysia.

^65^ Department of Urology, University of Washington, Seattle, WA, USA.

^66^ Institute of Human Genetics, University Medical Center Hamburg-Eppendorf, Hamburg, Germany.

^67^ Molecular Medicine Center, Department of Medical Chemistry and Biochemistry, Medical University, Sofia, Bulgaria.

^68^ Department of Oncology, Cross Cancer Institute, University of Alberta, Edmonton, Alberta, Canada.

^69^ Division of Radiation Oncology, Cross Cancer Institute, Edmonton, Alberta, Canada.

^70^ Molecular Endocrinology Laboratory, Department of Cellular and Molecular Medicine, KU Leuven, Leuven, Belgium.

^71^ Institute of Cancer Sciences, Manchester Cancer Research Centre, University of Manchester, Manchester Academic Health Science Centre, St Mary's Hospital, Manchester, UK.

^72^ Genomic Medicine Group, Galician Foundation of Genomic Medicine, Instituto de Investigacion Sanitaria de Santiago de Compostela (IDIS), Complejo Hospitalario Universitario de Santiago, Servicio Galego de Saúde, SERGAS, Santiago De Compostela, Spain.

^73^ University of California San Diego, Moores Cancer Center, La Jolla, CA, USA.

^74^ Department of Urology, Erasmus University Medical Center, Rotterdam, the Netherlands.

^75^ Cancer & Environment Group, Center for Research in Epidemiology and Population Health (CESP), INSERM, University Paris-Sud, University Paris-Saclay, Villejuif, France.

^76^ Clinical Gerontology Unit, University of Cambridge, Cambridge, UK.

^77^ Division of Genetic Epidemiology, Department of Medicine, University of Utah School of Medicine, Salt Lake City, Utah, USA.

^78^ George E. Wahlen Department of Veterans Affairs Medical Center, Salt Lake City, UT, USA.

^79^ The University of Surrey, Guildford, Surrey, UK.

^80^ Department of Laboratory Medicine and Pathology, Mayo Clinic, Rochester, MN, USA.

**Funding Acknowledgements**

Genotyping of the OncoArray was funded by the US National Institutes of Health (NIH) [U19 CA 148537 for ELucidating Loci Involved in Prostate cancer SuscEptibility (ELLIPSE) project and X01HG007492 to the Center for Inherited Disease Research (CIDR) under contract number HHSN268201200008I]. Additional analytic support was provided by NIH NCI U01 CA188392 (PI: Schumacher).

The PRACTICAL consortium was supported by Cancer Research UK Grants C5047/A7357, C1287/A10118, C1287/A16563, C5047/A3354, C5047/A10692, C16913/A6135, European Commission's Seventh Framework Programme grant agreement n° 223175 (HEALTH-F2-2009-223175), and The National Institute of Health (NIH) Cancer Post-Cancer GWAS initiative grant: No. 1 U19 CA 148537-01 (the GAME-ON initiative).

We would also like to thank the following for funding support: The Institute of Cancer Research and The Everyman Campaign, The Prostate Cancer Research Foundation, Prostate Research Campaign UK (now Prostate Action), The Orchid Cancer Appeal, The National Cancer Research Network UK, The National Cancer Research Institute (NCRI) UK. We are grateful for support of NIHR funding to the NIHR Biomedical Research Centre at The Institute of Cancer Research and The Royal Marsden NHS Foundation Trust.

**SUPPLEMENTARY FIGURES**

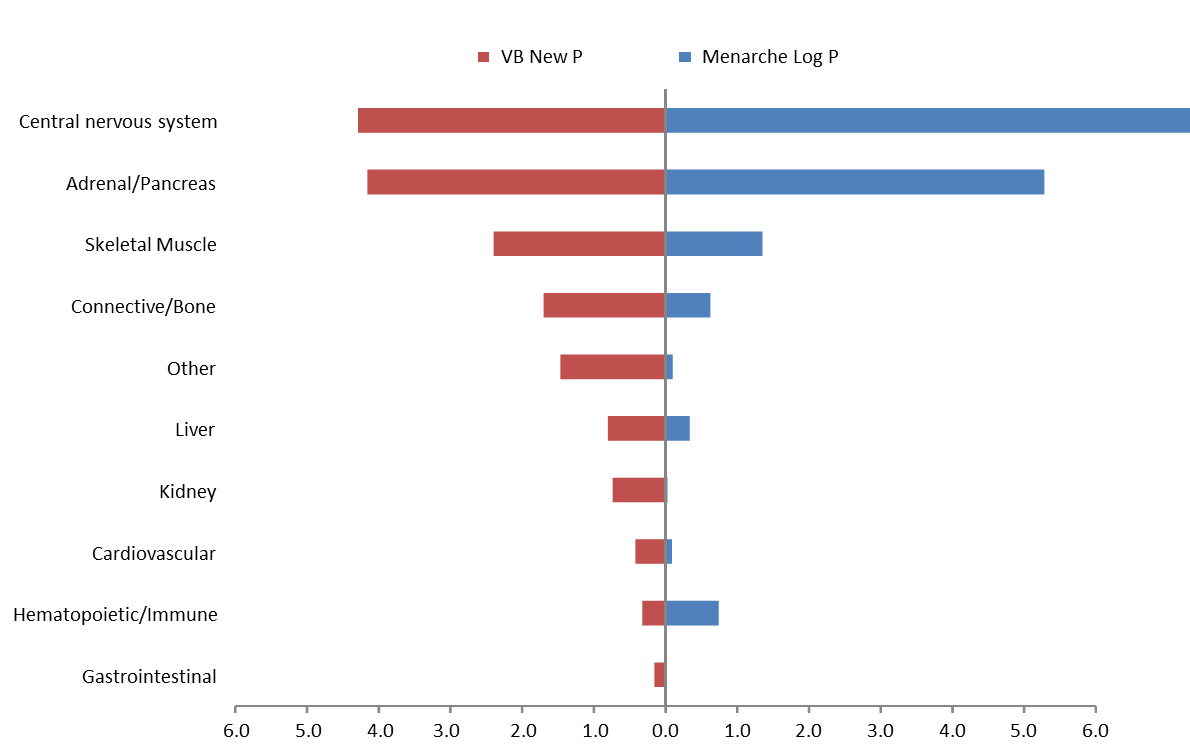

**Supplementary Figure 1:** Tissue expression (GTEx) for male puberty loci by general tissue categories.

-log10(P value)

|  | Adipose |
| --- | --- |
| Adipose (Subcutaneous) | 0.0761963 |
| Adipose (Visceral - omentum) | 0.173649 |
| Spleen | 0.102849 |
| Transformed lymphocytes | 0.051201 |
| Whole Blood | 0.4223 |
| Artery (Aorta) | 0.0791299 |
| Artery (Coronary) | 0.348674 |
| Artery (Tibial) | 0.381159 |
| Heart (Atrial Appendage) | 0.050828 |
| Heart (Left Ventricle) | 0.223459 |
| Adrenal Gland | 1.27931 |
| Ovary | 0.878766 |
| Pancreas | 0.919992 |
| Testis | 0.270898 |
| Thyroid | 0.822956 |
| Colon (Sigmoid) | 0.648776 |
| Colon (Transverse) | 0.0478627 |
| Esophagus (Gastroesophageal Junction) | 0.539669 |
| Esophagus (Mucosa) | 0.00982896 |
| Esophagus (Muscularis) | 0.554409 |
| Small Intestine (Terminal Ileum) | 0.0369345 |
| Stomach | 0.702736 |
| Skeletal Muscle | 1.3385 |
| Brain (Amygdala) | 0.705268 |
| Brain (Anterior cingulate cortex) | 1.42397 |
| Brain (Caudate) | 0.855511 |
| Brain (Cerbellum) | 1.85031 |
| Brain (Cerebellar Hemisphere) | 2.15882 |
| Brain (Cortex) | 1.96047 |
| Brain (Frontal cortex) | 1.51445 |
| Brain (Hippocampus) | 0.425054 |
| Brain (Hypothalamus) | 1.92456 |
| Brain (Nucleus accumbens) | 0.932996 |
| Brain (Putamen) | 1.04227 |
| Brain (Substantia nigra) | 0.588376 |
| Cervical Spinal Cord | 0.840239 |
| Nerve (Tibial) | 0.30669 |
| Pituitary | 1.16853 |
| Bladder | 0.227071 |
| Breast (Mammary) | 0.406851 |
| Cervix (Ectocervix) | 0.0354849 |
| Cervix (Endocervix) | 0.129333 |
| Fallopian Tube | 0.226189 |
| Kidney (Cortex) | 0.452931 |
| Liver | 0.682309 |
| Lung | 0.168933 |
| Minor Salivary Gland | 0.42765 |
| Prostate | 1.20443 |
| Skin (Not Sun Exposed - Suprapubic) | 0.00958006 |
| Skin (Sun Exposed - Lower leg) | 0.0237748 |
| Transformed fibroblasts | 0.333039 |
| Uterus | 1.09923 |
| Vagina | 0.0801872 |

**Supplementary Figure 2:** Tissue expression for male puberty loci by specific GTEx tissues.

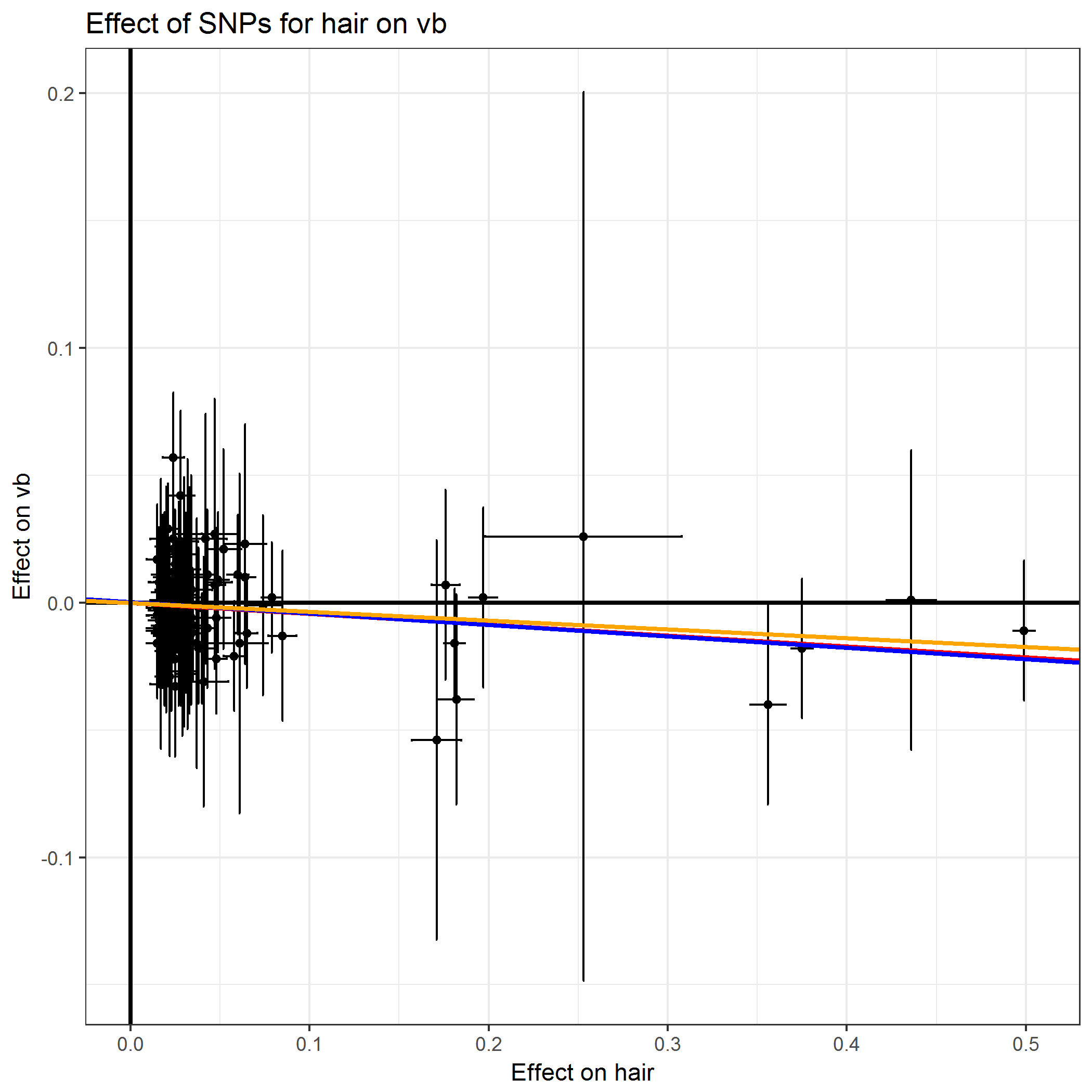

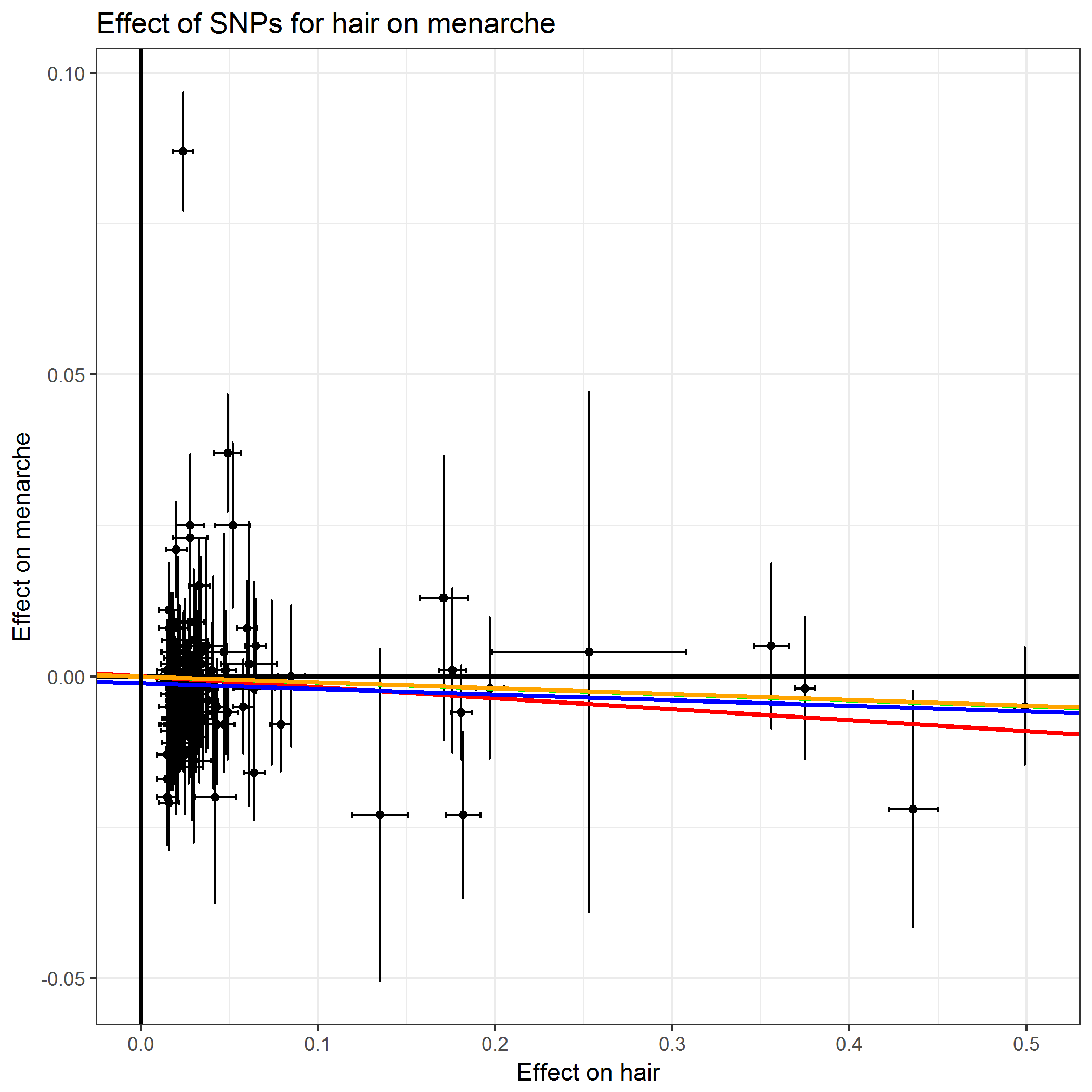

**Supplementary Figure 3:** Mendelian randomisation analyses for effect of hair pigmentation on puberty timing in men (left panel) and women (right panel).

|  | rg |
| --- | --- |
| Body mass index (BMI) | -0.3214 |
| Pain all over the body | -0.2821 |
| Long-standing illness, disability or infirmity | -0.248 |
| Osteoarthritis (self-reported) | -0.2405 |
| Body fat percentage | -0.2335 |
| Hip pain | -0.2259 |
| Waist-hip ratio | -0.2108 |
| Smoking (pack-years) | -0.2042 |
| Diabetes | -0.2034 |
| Alcohol intake (frequency) | -0.203 |
| Angina | -0.2013 |
| Number of treatments/medications | -0.1967 |
| Number of non-cancer illnesses | -0.175 |
| Back pain | -0.1662 |
| Hypertension (doctor-diagnosed) | -0.1605 |
| Heavy manual or physical work | -0.1604 |
| Systolic blood pressure | -0.1305 |
| Fluid intelligence | 0.1332 |
| Education: College or University degree | 0.1784 |
| Overall health rating | 0.22 |
| Age at first live birth | 0.2306 |
| Usual walking pace | 0.2579 |
| Mother's age at death | 0.2662 |
| Age at menarche (females) | 0.6827 |
| Adult height minus height at age 14 | 0.7729 |

**Supplementary Figure 4:** Genetic correlations (r_g_) between male puberty timing and selected anthropometric and health-related traits, calculated using LD Score regression. Positive correlations are shown in blue and negative correlations in red. Error bars represent 95% confidence intervals.

**a)**

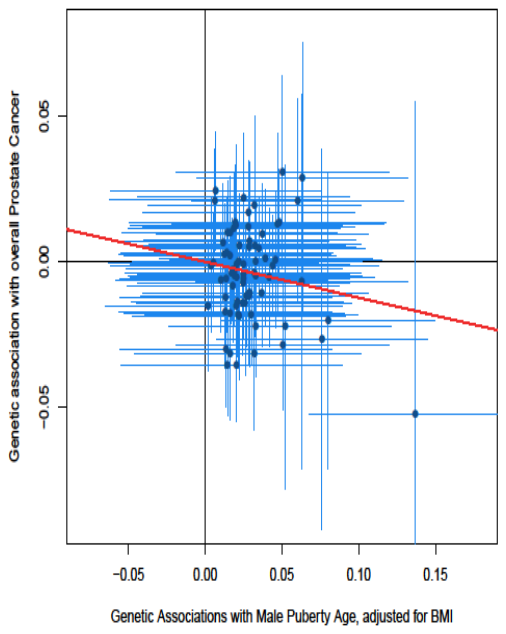

**b)**

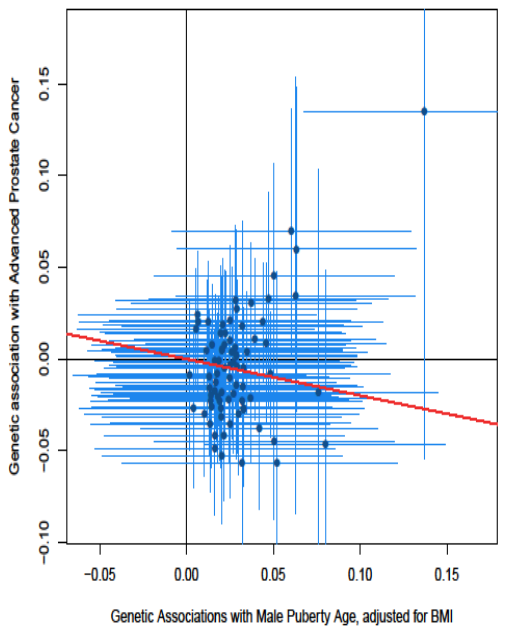

**Supplementary Figure 5:** Mendelian randomisation analysis (random effects IVW method) for puberty timing (adjusted for BMI) to a) overall prostate cancer risk and b) risk of advanced prostate cancer.
